# Quantitative Modeling and Automated Analysis of Meiotic Recombination

**DOI:** 10.1101/083808

**Authors:** Martin A. White, Shunxin Wang, Liangran Zhang, Nancy Kleckner

## Abstract

Many morphological features, in both physical and biological systems, exhibit spatial patterns that are specifically characterized by a tendency to occur with even spacing (in one, two or three dimensions). The positions of crossover (CO) recombination events along meiotic chromosomes provides an interesting biological example of such an effect (1–3). In general, mechanisms that explain such patterns may (a) be mechanically-based, (b) occur by a reaction-diffusion mechanism in which macroscopic mechanical effects are irrelevant, or (c) involve a combination of both types of effects. We have proposed that meiotic CO patterns arise by a mechanical mechanism, have developed mathematical expressions for such a process based on a particular physical system with analogous properties (the so-called “beam-film model”), and have shown that the beam-film model can very accurately explain experimental CO patterns as a function of the values of specific defined parameters (4–7). Importantly, the mathematical expressions of the beam-film model can apply quite generally to any mechanism, whether it involves mechanical components or not, as long as its logic and component features correspond to those of the beam-film system (3; below). Furthermore, via its various parameters, the beam-film model discretizes the patterning process into specific components. Thus, the model can be used to explore the theoretically predicted effects of various types of changes in the patterning process. Such predictions can expand detailed understanding of the bases for various biological effects (e.g. 2, 5). We present here a new MATLAB program that implements the mathematical expressions of the beam-film model with increased robustness and accessibility as compared to programs presented previously. As in previous versions, the presented program permits both: (i) simulation of predicted CO positions along chromosomes of a test population; and (ii) easy analysis of CO positions, both for experimental data sets and for data sets resulting from simulations. The goal of the current presentation is to make these approaches more readily accessible to a wider audience of researchers. Also, the program is easily modified, and we encourage interested users to make changes to suit their specific needs. A link to the program is available on the Kleckner laboratory web site: http://projects.iq.harvard.edu/kleckner_lab.

## 1. Introduction

Meiosis is the specialized cellular program that underlies halving of the chromosome complement (e.g. from diploid to haploid) as required for gamete formation and sexual reproduction. A central component of meiosis is recombination, which plays both evolutionary and mechanistic roles (1, 2). During this process, a large number of recombinational interactions are initiated via programmed DNA double-strand breaks (DSBs). Most DSBs identify and engage the corresponding region on a homologous chromosome (i.e. the maternal or paternal “homolog”). These inter-homolog interactions, via their association with chromosome structure components, concomitantly mediate whole chromosome pairing, with coalignment of homologs along their lengths.

At about this coalignment stage (discussions in 2, 7), a small subset of these total interhomolog interactions are specifically designated for eventual maturation into crossover (CO) products, where flanking regions on the two involved chromatids are reciprocally exchanged. These specifically designated CO interactions occur at different positions in different meiotic nuclei; nonetheless, they tend to be evenly spaced. This pattern was originally identified as the classical phenomenon of CO interference: the frequency of occurrence of a CO at one position along a chromosome is reduced if that chromosome also exhibits another CO nearby. Interhomolog recombinational interactions that are not designated to become COs via this patterning process will, instead, mature to other fates.

We have proposed that CO patterning occurs by a stress-and-stress-relief mechanism (4; Fig. 1). In brief, all early (undifferentiated) inter-homolog interactions come under mechanical stress, which finally begins to promote CO-designation events. We refer to the interactions upon which CO-designation acts as “precursors”. When stress-promoted CO-designation occurs at a particular position, it will necessarily involve molecular changes that send the affected interaction down the CO pathway and, concomitantly, will result in local alleviation of stress at the affected site. The mechanical nature of the system implies that this change in stress will redistribute, moving outward from its nucleation site and tending to even out the level of stress along the length of the chromosome. However, this effect will tend to be absorbed by chromosomal components as it spreads and thus will tend to dissipate with distance. The consequence will be a self-limiting zone of reduced stress, i.e. a zone of “CO interference”, within which the probability of a subsequent CO-designation is commensurately reduced. Any second stress-promoted CO-designation will tend to occur outside of this first zone, where the stress level remains high, and will create a second zone of interference. Subsequent CO-designations will tend to occur away from the positions of prior designations (and their interference zones), “filling in the holes” between previous events and ultimately giving even spacing. Implicit in this description is the fact that, at any given moment in the patterning process, each “precursor” will have a potential to undergo CO-designation which is determined by (the product of) the intrinsic sensitivity of that precursor to stress and the local level of stress at that position at that point in time, which may or may not have been effected by stress relief (interference) emanating across that position from a nearby CO-designation.

**Fig. 1.**
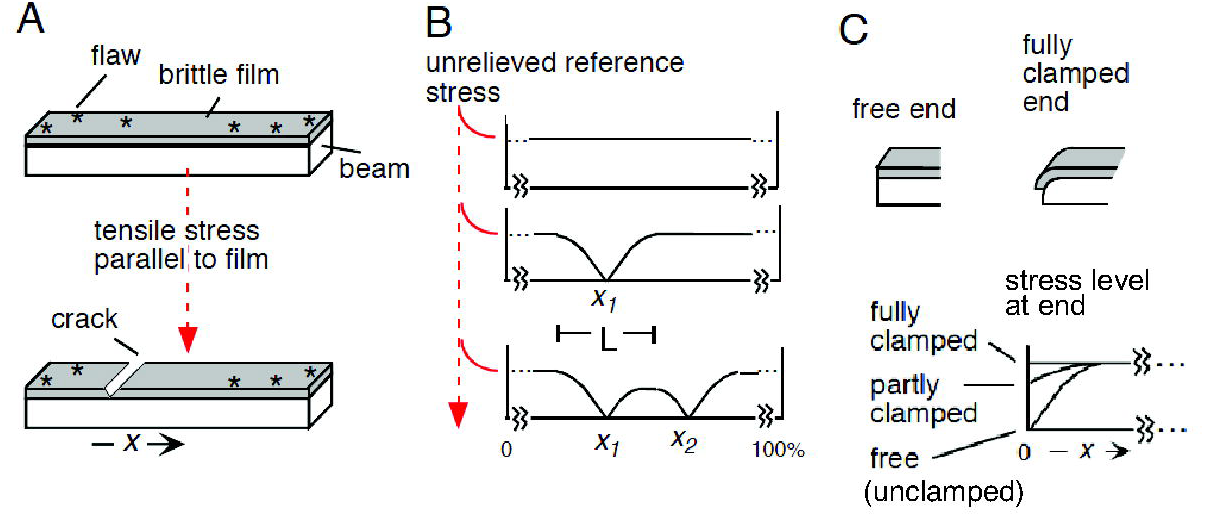
The beam-film model for meiotic CO patterning. (**A**) An elastic beam coated with a thin, brittle film containing flaws experiences stress along the beam/film interface (e.g. from differential expansion of the beam in response to temperature). Stress will promote cracking of the film at flaw sites. The beam/film is analogous to a meiotic prophase bivalent; flaws are analogous to DSB-mediated “precursors”; and cracks are analogous to COs. (**B**) Sequential occurrence of cracks progressively alters the level of stress along the beam (in both directions) over a characteristic length (L). Stress-promoted occurrence of cracks is disfavored in regions of lower stress. Sequential cracks tend to fill in the holes between the zones of stress relief of preexisting cracks. (**C**) The ends of the beam/film ensemble may be “clamped”, with stress well supported at the end; “free” (“unclamped”), or partially clamped. At a free end, the level of stress at a free end falls to zero, giving the same effect as a pre-existing crack. Partial clamping results in an intermediate effect. (Reproduced from (4) with permission, Copyright (2004) National Academy of Sciences, U.S.A.).

The above description makes it clear that the same effects could arise in many ways as long as there is a set of initial “precursor” interactions and a patterning process that involves: (i) a tendency for CO-designation; (ii) an intrinsic sensitivity of each precursor to that CO-designation tendency; and (iii) an effect in which local CO-designation nucleates formation of a signal that is inhibitory to CO-designation, spreads outward in both directions from the nucleating site, and dissipates with distance.

The final observed array of meiotic COs also depends upon effects that occur after CO-designation. First, a CO-designated interaction must undergo a multiplicity of ensuing biochemical steps in order to finally become a CO product. We refer to this process as “CO maturation". Second, occasionally, an interaction that has not been designated to be a CO as part of the patterning process will, nonetheless, produce a CO product. Such products are referred to as “Type II” COs (1) and are detected by some experimental assays but not by others (e.g. 8).

One physical system that exhibits stress-and-stress relief effects analogous to those described above is an elastic beam coated with a thin brittle film that contains flaws. Stress along the beam/film interface causes a flaw(s) to become a crack(s) that extend across the beam perpendicular to its length, thereby alleviating stress along the beam to either side of the crack, to decreasing extent with increasing distance. Within this zone of stress relief, the probability of formation of subsequent stress-promoted crack(s) is reduced, in relation to the magnitude of stress relief. The beam-film ensemble corresponds to a prophase chromosome ("bivalent"). Flaws are analogous to precursor recombinational interactions. A crack is analogous to a CO and the resulting local domain of stress relief is analogous to a zone of CO interference. We have previously presented mathematical expressions, implemented by appropriate software, that enable modeling of CO-formation according by this beam-film scenario and, by extension, any other process that works in the analogous way.

For purposes of such modeling, CO patterning is divided into three aspects, each of which is appropriately parameterized: (I) the array of precursor interactions; (II) the nature of the patterning process itself; and (III) the effects of post-patterning events. Discretization of CO patterning into these different parameterized aspects makes it possible to begin to think in more mechanistic detail about how the process could work. To this end, expressions of the beam-film model make it possible to simulate the number and pattern of COs that are predicted to occur under any specific set of values of the involved parameters. Such simulations can be used in two ways. First, given an experimental data set, it is possible to determine the set of parameter values whose predicted outcome best matches the data. Such “best-fit simulations", as performed for a number of organisms, including wild type and mutant situations, show that the “beam-film” model can very accurately describe experimental data and also can provide a framework for understanding the effects of mutations and other genetic variations. Second, the beam-film model can be used to explore the effects of, and interplay among, different aspects of the patterning process in a theoretical sense, thereby deepening our detailed understanding of potential effects, generating new hypotheses, and motivating quantitative analyses.

We present here an updated version of our previously published MATLAB program for beam-film simulations along with detailed instructions for its use. The current version is improved with respect to both robustness and accessibility. In addition, this version enables automated analysis of CO distributions provided either by experimental data or as the outputs of beam-film simulations.

## 2. Materials

1. Copy of the MATLAB program ‘Crossover Patterning Simulation and Analysis 1.0’. A folder (named ‘Crossover_Simulation_and_Analysis’) containing the 16 MATLAB files of the program is available for download *via* the authors lab website http://projects.iq.harvard.edu/kleckner_lab (*see* **Note 1**).
2. Computer with MATLAB software (*see* **Note 2**).

## 3. Methods

### 3.1 Terminology: Bivalents Versus Chromosomes

Meiotic COs link maternal and paternal homologs from prophase through metaphase I. Each such pair is referred to as a “bivalent” because of this dual nature. The term “chromosome” will be used here in the sense of its genetic identity, e.g. “chromosome 21", rather than in the sense of a physical object. Thus, in an individual nucleus, COs occur along a given bivalent which corresponds to a particular genetic chromosome. In a population of meiotic nuclei, CO patterns for a particular chromosome are defined by analysis of the many bivalents that occur in the corresponding many nuclei. In a beam-film simulation analysis, CO positions are defined along each of a specified number of bivalents, thus representing the positions along the bivalent corresponding to a particular chromosome in a corresponding number of different nuclei.

### 3.2 Descriptions of Parameters

A given beam-film simulation requires the user to specify the values of all parameters in the three categories outlined above.

#### 3.2.1 Group I: Precursor Parameters

##### N, B and E

The array of undifferentiated precursors upon which CO patterning acts is defined by three basic parameters: N, B and E. **N =** the average number of precursors per bivalent. **B =** the extent to which the number of precursors per bivalent varies from one nucleus to another. The value of B ranges from 0 to 1. 0 specifies maximum variation as described by a Poisson distribution with a mean value of (N); 1 specifies absence of variation, with the same constant number of (N) precursors on the bivalent in question in all nuclei; intermediate values are defined by appropriate binomial distributions (e.g. Fig. 2A). Thus: N and B together describe the number of precursors on each bivalent in the set of bivalents to be analyzed. **E =** the extent to which the precursors along each given bivalent are randomly versus evenly spaced (Fig. 2B). The value of E ranges from 0 to 1, analogously to B. For E=0, positions are drawn at random from a standard uniform distribution. E=1 represents perfectly even spacing.

**Fig. 2.**
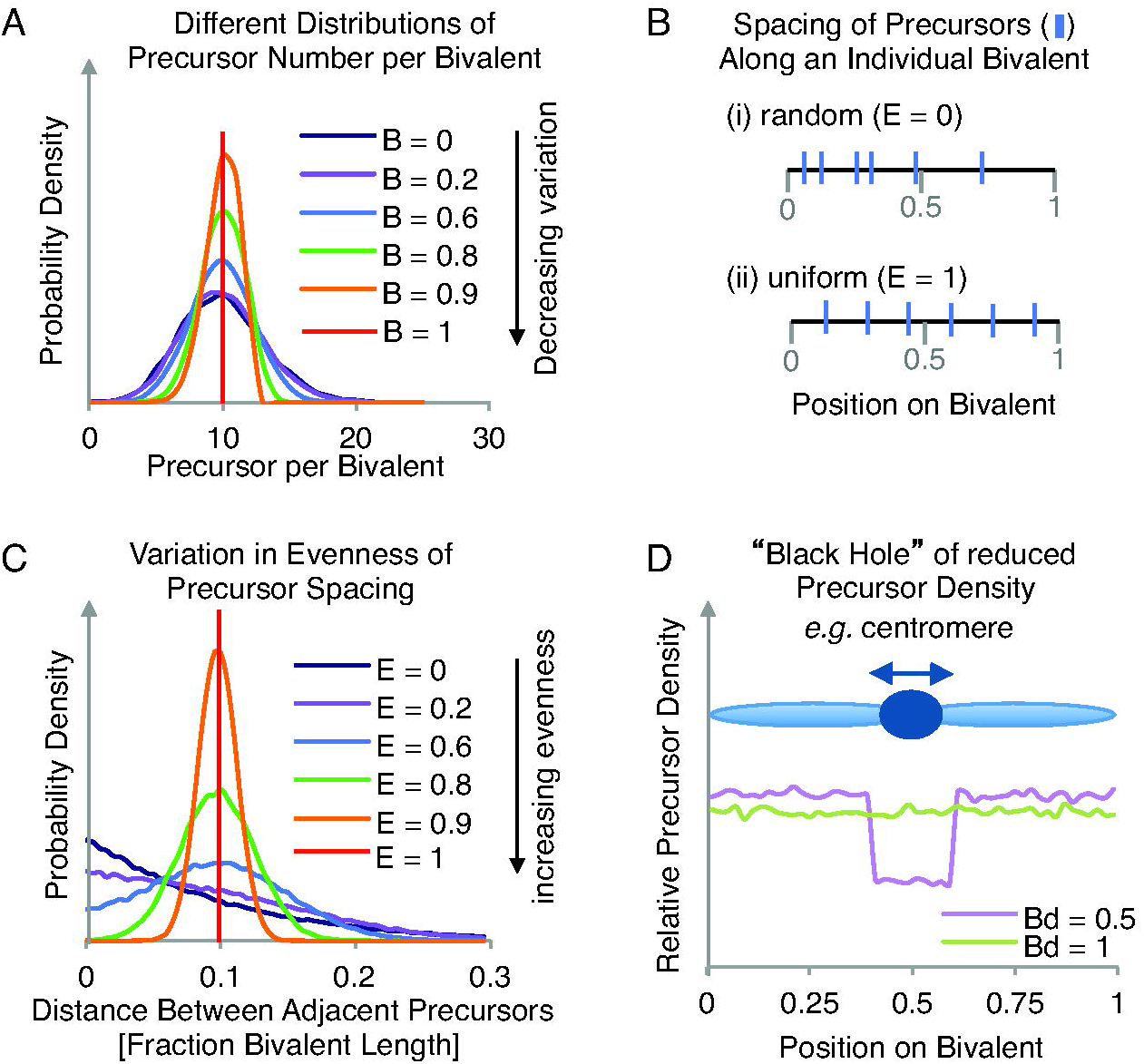
Creating an array of precursors on each simulated bivalent. (**A**) The effect of varying parameter B on the distribution of total precursor numbers (when N=10 and E=0). (**B**) The qualitative effect of increasing parameter E from (**i**) 0 to (**ii**) 1 is to make the precursors more evenly spaced along each simulated bivalent. (**C**) The effect of varying parameter E on the distribution of distances between adjacent precursors (when N=10 and B=0). (**D**) The effect of varying parameter Bd on the relative precursor density along the bivalent length (when Bs=0.4 and Be=0.6). Simulating a ‘Black Hole’ is useful for modeling events across, for example, centromeres (dark blue circle on light blue ‘bivalent’).

Intermediate values representing incomplete tendencies for even spacing. More and less even spacing results in narrower and broader distributions of distances between adjacent precursors (Fig. 2C). In general, there is a tendency for precursor interactions to be evenly spaced along meiotic chromosomes and for the number of precursors per bivalent to be relatively (but not perfectly) constant (e.g. discussions in 5, 7, 9, 10 and Zhang unpublished).

##### Black hole: Bs, Be and Bd

The program also offers the possibility of creating a “black hole", i.e. a region in which the average number of precursors is less than the density along the rest of the chromosome. This feature is useful because the frequency of recombination-initiating DSBs is known to be dramatically reduced in centromeric regions, with commensurate reductions in the frequencies of COs in these regions. An example of a black hole pattern is shown in Fig. 2D. The nature of the black hole is specified by three parameters: **Bs** (the start of the region of precursor suppression), **Be** (the end of the region of precursor suppression) and **Bd** (the precursor density within this region relative to the density elsewhere along the bivalent). Given values for Bs and Be, the black hole is implemented programmatically by: (i) selecting a number of precursors (parameter N) corresponding to that expected for the desired frequency along the entire bivalent in the case where the black hole would be absent; (ii) distributing those precursors according to parameters B and E; and (iii) considering each precursor in the black hole region and removing it with a probability defined by the value of Bd. We note that the program can be readily modified by an interested user to include more than one black hole.

#### 3.2.2 Group II: Patterning Parameters

Patterning of CO designation events is best described in the context of the beam-film stress hypothesis; however, it must be kept in mind that all of the patterning parameters have generic analogues that would pertain analogously to any mechanism.

##### Smax, A and L

Patterning is defined by three basic parameters: Smax, A and L. The potential of a particular precursor interaction to undergo CO-designation at some particular moment during the process is given by the product of two parameters: (intrinsic sensitivity of the precursor to stress) x (the level of stress present at the corresponding position), a feature we refer to as “Local Crossover Potential” or “LCP” (this value equates to variable “strc” in the program code). At each step, the precursor that undergoes CO-designation is the one with the highest LCP; and the CO-designation process continues until there is no remaining precursor for which LCP > 1. The process thus proceeds as follows (assuming that the ends are considered to be “clamped", as described below). Prior to the first CO-designation, the level of stress is at a particular “starting level” all along the bivalent as defined by the parameter **Smax** and each precursor has its own individual sensitivity to stress, which is defined by implementation of parameter **A** as described below. The first CO-designation will occur at the site of the most sensitive precursor (which will have the highest LCP). This will cause the level of stress to fall to zero at the site of the CO-designation and will nucleate a zone of reduced stress that spreads outwards in both directions from that position, dissipating exponentially with distance (a.k.a. interference). The characteristic distance over which stress redistributes (in both directions, Fig. 1B) is defined by the parameter **L**, also known as the “stress relief distance” or “interference distance". Next, the program recalculates the LCPs for all remaining precursors, taking into account the changes in stress levels resulting from the CO-designation, after which a next CO-designation occurs, again at the position of the precursor with the highest LCP, again triggering changes in the level of stress in the surrounding region. These steps of LCP recalculation, CO-designation and spreading interference are repeated until no remaining precursor has an LCP > 1.

In a more general model for CO interference: (i) **Smax** would represent the strength of any CO-designation process; (ii) implementation of **A** would allow the definition of intrinsic sensitivities of precursors to that process; and (iii) interference could result from a decrease in the sensitivities of affected precursors to a CO-designation process of constant strength, with spreading and dissipation with distance as described by the parameter **L**.

##### Intrinsic precursor sensitivities (A)

The intrinsic sensitivities of precursors to stress are determined as follows. Each precursor is assigned a number from 0 to 1 from a uniform distribution of total precursors. This assignment ensures that, along a given bivalent, every precursor will have a different sensitivity from every other precursor. Moreover, because of the uniform distribution, every precursor sensitivity value is as probable as every other precursor sensitivity value. The sensitivity levels defined in this way are the default option in the program, which is specified by a value of A=1. This procedure ensures that CO-designations along a given bivalent occur sequentially and also defines a particular range and distribution of precursor sensitivity values.

However: it has also been useful to consider distributions of precursor sensitivities other than that provided by a uniform distribution ranging from 0 to 1. For this purpose, the values assigned by A=1 can be transformed into another set of values using any one of several nonlinear functions. Each such function is represented in the program by a different value of A (currently A=2 to A=7). Each such transformation again yields an array of equally probable precursor sensitivities; however, the absolute and relative values of these sensitivities are different in each case, as specified by the corresponding function (Fig. 3).

**Fig. 3.**
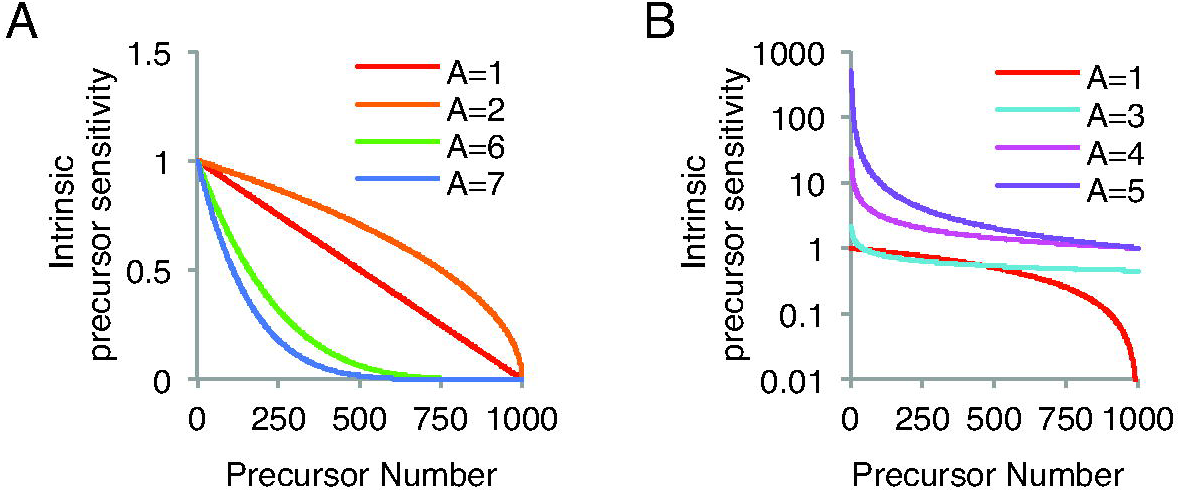
The effect of varying parameter A on the distribution of intrinsic sensitivity values for 1000 simulated precursors. Note that the intrinsic precursor sensitivities in panel (**A**) are plotted on a linear scale and the intrinsic precursor sensitivities in panel (**B**) are plotted on a logarithmic scale.

##### Bsmax

Parameter **Bsmax** makes it possible for different bivalents in a population to have different levels of Smax. Values of Bsmax vary from 0 to 1 where, analogously to B (above), 0 is a Poisson distribution about Smax (μ = Smax), 1 is a constant value of Smax for all bivalents, and intermediate values are appropriate binomial distributions.

##### cL and cR

Parameters **cL** and **cR** define what happens at the left and right end of a bivalent respectively. In a true beam-film system (Fig. 1C), an end may be completely free. In this case, stress is not supported at the end, which thus behaves as a “pre-existing crack": it is as if there is already a CO at the end of the bivalent even before the first CO-designation event. At the other extreme, stress is fully supported at the end, e.g. by wrapping of the film around the end of the beam. In this case, the level of stress present prior to the first CO-designation is the same at the end as elsewhere along the beam. Moreover, interference necessarily cannot emanate into a terminal region from “beyond the end of the bivalent". Thus, in this case, the frequency of COs will be higher at the end than for an internal region. In these two situations, the end is said to be either “unclamped” or “clamped", respectively. Intermediate levels of clamping are also possible. These conditions are described at the left and right ends of the bivalent by the parameters cL and cR respectively, where cL/cR varies continuously from 0 (unclamped) to 1 (clamped). It is possible to envision a direct analog of the clamped state: in many organisms, chromosome ends are robustly attached to the nuclear envelope at the time of CO-designation. More generally, however, variations in cL/cR can be used to model “end effects".

#### 3.2.3 Group III: Post-patterning Parameters

##### M

The multiple additional steps required for maturation of a CO-designated recombinational interaction into a final CO product may occur efficiently or not. Variations in maturation efficiency are described by parameter **M**, which varies from 0 (no maturation) to 1 (100% maturation). A paradigmatic example of maturation inefficiency is provided by elimination of MutL homolog Mlh1 (5).

##### T2prob

The contribution of Type II COs appear to arise as a low probability outcome at non-CO-designated sites (Introduction) and can be taken into account in the final CO output using the parameter **T2prob**, which is implemented after CO-designation process is complete. This parameter defines, for each site where CO-designation has not occurred, the probability that a Type II crossover will occur. The value of T2prob can vary from 0 to 1.

### 3.3 Describing CO Patterns

Experimental analysis provides a set of data comprising the positions of COs along each of a set of bivalents corresponding to events along a particular chromosome in a corresponding set of nuclei. Beam-film simulations provide an exactly analogous set of data, with the exception that multiple independent data sets, each comprising very large numbers of “bivalents” (usually 5000 - 10,000), are easily obtained for any given set of parameter values.

#### Automatic analyses

Both types of outputs can be analyzed, analogously, as desired. Towards this end, the beam-film program automatically carries out several commonly-used analyses, providing the following information: (i) the average number of COs per bivalent; (ii) the distribution of frequencies of bivalents exhibiting different numbers of COs (0, 1, 2…etc); (iii) the density distribution of CO frequencies along the length of the analyzed chromosome; (iv) Coefficient of Coincidence (CoC) relationships (further discussion below); (v) the average distance between adjacent COs; (vi) the distribution of distances between adjacent COs; and (vii) fitting of the distribution of distances between adjacent COs by a gamma distribution, where the shape parameter (*Κ*) of the best-fit distribution is taken as a measure of “evenness” of inter-CO distances and thus CO spacing.

#### CoC Analysis

CoC analysis provides the clearest picture of CO patterning because it directly reports communication along individual bivalents. In contrast, as discussed previously (5), the gamma distribution shape parameter can change because of features unrelated to CO patterning, e.g. changes in maturation efficiency.

CoC analysis is performed as follows (further details in 5): (i) the chromosome of interest is divided into intervals, which may be of equal or unequal size as desired; (ii) CO frequencies are defined for each interval individually; (iii) intervals are then considered in all possible pairwise combinations with respect to (a) the frequency of “double COs” in the data set, i. e. the frequency of bivalents that have a CO in each of the two intervals of a pair, and (b) the frequency of double COs expected if COs occurred independently in the two intervals, given by the product of the frequencies in each interval considered individually (above). The ratio of “observed” to “expected” double COs is the Coefficient of Coincidence (CoC). CoC values for all pairs of intervals are plotted as a function of inter-interval distance (averaging the values for all pairs separated by the same distance if appropriate). In the classical outcome (e.g. Fig. 4A), the CoC is low (or zero) for short inter-interval distances (reflecting strong interference) rises with increasing inter-interval distance to a value of 1 (reflecting decreasing interference with increasing inter-interval distance until the point where events in two intervals are independent) and then a tendency to fluctuate above 1 at periodic intervals (reflecting the tendency for COs to be evenly spaced along each bivalent with a certain periodicity). The nature of CO patterning is described by the CoC curve as it rises from small inter-interval distances to a value of 1. A convenient metric to describe this feature is the inter-interval distance at which CoC = 0.5, a value we define as L_CoC_ (5; Fig. 4A).

**Fig. 4.**
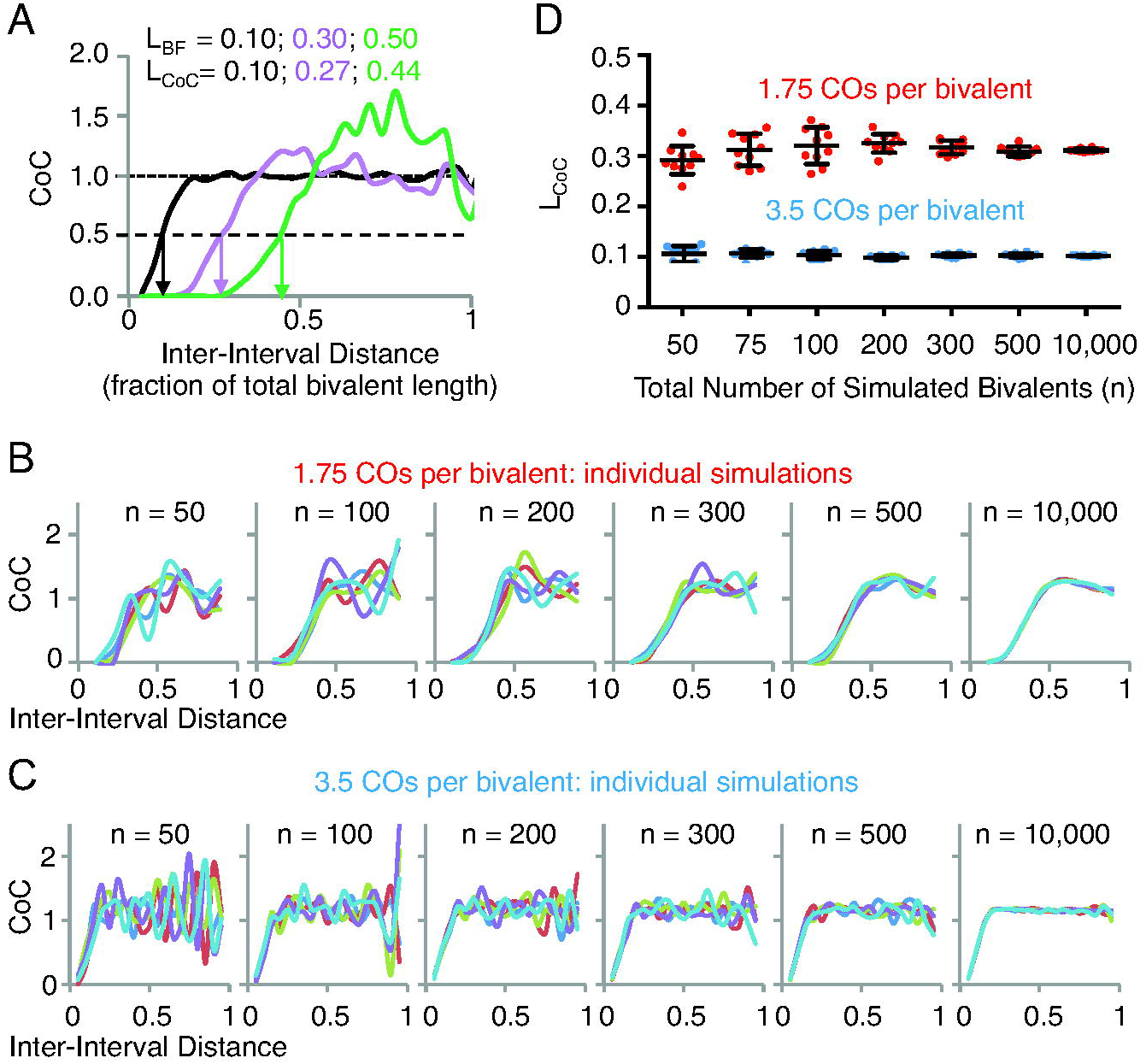
Sample size and coefficient of coincidence (CoC) analysis. (**A**) The effect of varying parameter L (L_BF_) from 0.1 to 0.5 on CoC curves and the inter-interval distance at which the CoC value is 0.5 (L_CoC_). (**B-D**) The effect of varying parameter n (the total number of simulated bivalents) on the reproducibility of CoC analysis using parameters that gives rise to an average of either 1.75 (red) or 3.5 (blue) crossovers per bivalent. Ten independent simulations were carried out for each value of n. (**B**) and (**C**) CoC curves for five of the ten independent simulations for n=50 to 10,000 bivalents. (**D**) The L_CoC_ values for each of the ten independent simulations for n=50 to 10,000). Each circle represents a data point. Bars indicate the mean and standard deviation.

As a practical matter, meaningful CoC analysis requires that the number of observed double COs be large enough to give an accurate set of CoC relationships. This is not a problem for data sets provided by simulation, where the number of bivalents can be as large as desired. However, it is a consideration for experimental data. In principle, the number of double COs will be a function of three interacting variables: (i) average number of COs per bivalent; (ii) sample size (number of bivalents analyzed); and (iii) number of intervals into which the chromosome is divided for analysis. The lower the average number of COs per bivalent, the larger the sample size required for reliable analysis; nonetheless, accurate evaluation can be achieved with as few as 100-200 bivalents, even with fewer than 2 COs per bivalent (Fig. 4B-D). Further, in general, we find that the interval size should be less than ~ 1/4 the average distance between adjacent COs. On the other hand, the smaller the interval size, the lower the frequency of double COs and thus, for a given average frequency of COs per bivalent, the larger the data set required.

### 3.4 Examples of Applications: Describing and Analyzing COs on Human Male Chromosomes (13-16)

The ability of the beam-film model to accurately describe experimental data, documented previously (5–7) can be further illustrated by applying the improved program presented here to best-fit simulation analysis of CO positions along human male meiosis chromosomes (13-16), which are similar enough to be considered as a group. CO positions are defined by CO-correlated Mlh1 foci along pachytene bivalents (11). Application of the program’s analysis function to experimental data illustrates resultant plots for: frequencies of bivalents with different numbers of COs, the CO probability density along the length of the bivalent, and CoC relationships (Fig. 5, black). Best-fit simulation analysis identifies the set of parameter values that provides the most accurate description of the experimental data, as evaluated by comparison of the same descriptors (Fig. 5, red; Table 1).

**Table 1.**
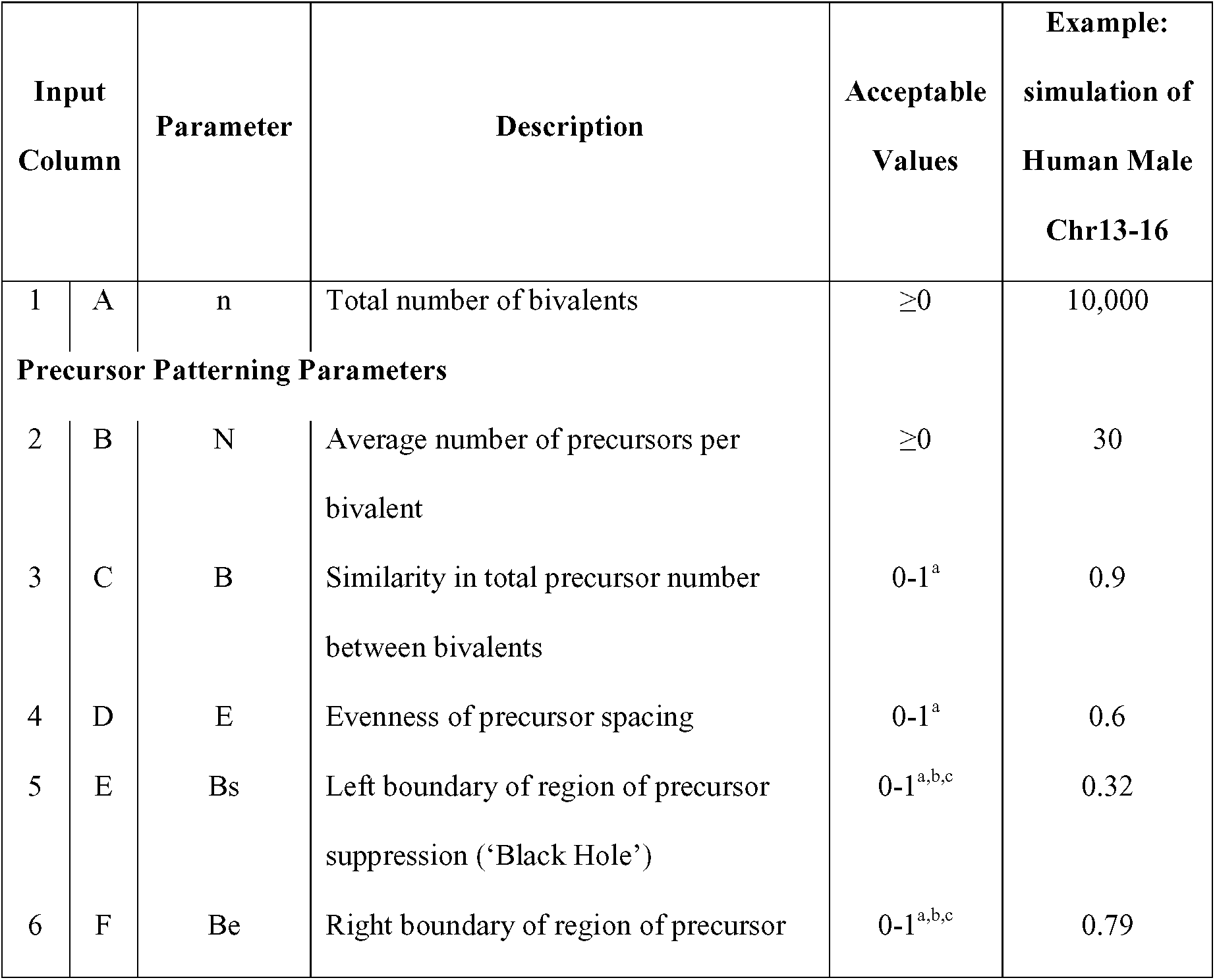
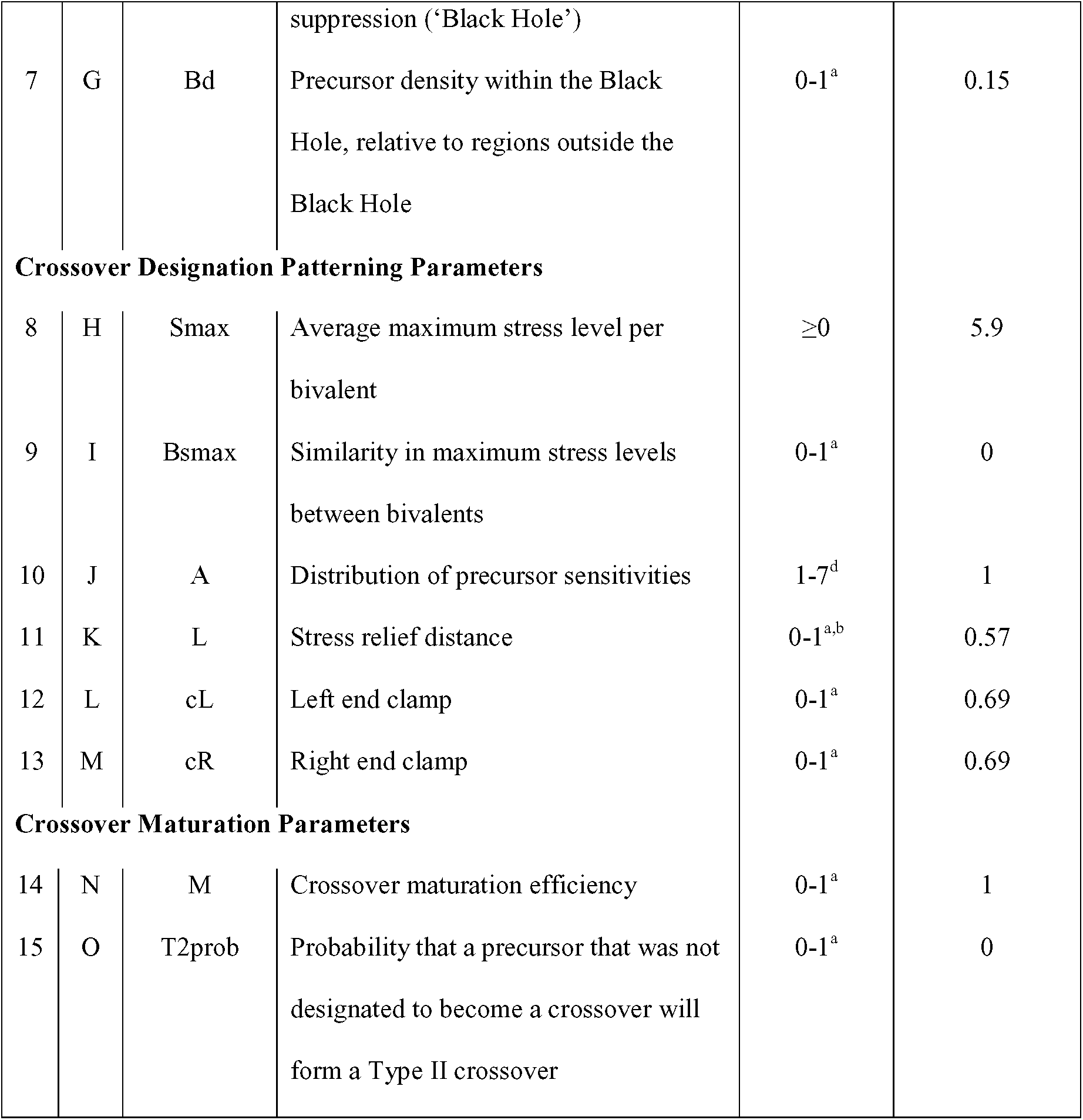
Specification of parameter values for beam-film simulations. Simulations require the user to input values for 15 parameters in the format of a table saved as a comma separated values (.csv) file. Values for different parameters are entered in different columns (as indicated as ‘Input Column’) and parameter values for separate simulations are entered on different rows. The table must contain column headers. Descriptions of the parameters, limits on their possible values and as an example, the parameter values used to simulate the distribution of Mlh1 foci on chromosomes 13-16 of human male meiosis chromosomes (Figure 5) are shown. An example input table is included with the program (simulation_parameters.csv). ^a^Values less than 0 are treated as 0, values greater than 1 are treated as 1. ^b^Values are relative to the full length of the bivalent (*i.e.* a fraction of 1). ^c^The maximum size of the Black Hole corresponds to Bs=0.001 and Be=0.999. ^d^Other values will result in A=1.

**Fig. 5.**
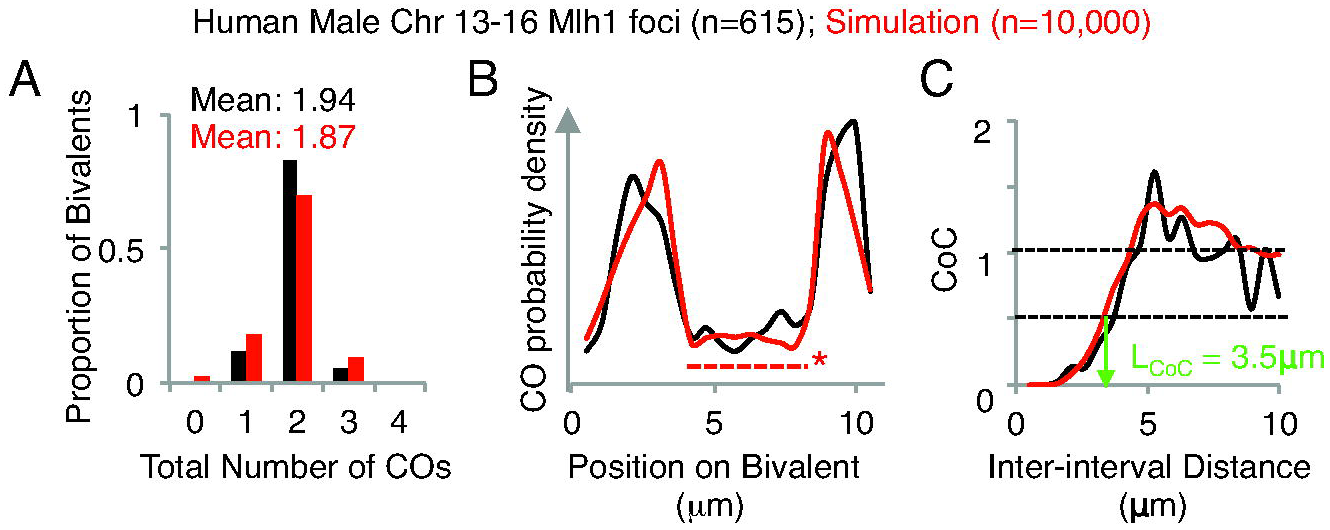
The distribution of Mlh1 foci along human male chromosomes 13,14, 15 and 16 were analyzed (615 bivalents in total) and simulated (10,000 bivalents total). Values for data are shown in black, values for simulation are shown in red. The parameters used in this simulation can be found in Table 1. (**A**) The mean number of total COs and the distribution of total CO numbers across the population. (**B**) The density distribution of COs along the length of chromosome(s). The dashed line with an asterisk indicates the region of the chromosome corresponding to the simulated ‘Black Hole’. (**C**) Coefficient of coincidence analysis. L_CoC_ 0.5 (the inter-interval distance at which the CoC value is 0.5) is approximately 3.5 μm for both the data and the simulation.

#### Quantitative analysis of the patterning process

The code can easily be modified to produce each intermediate value in the patterning process. For example, a bivalent with 21 evenly spaced precursors along the full length of the bivalent (n=1, N=21, Bs=0, Be=0, Bd=1) was simulated. These precursors were assigned intrinsic sensitivities in accord with parameter A=1. The pattern_event_designations_according_to_beam_film_model function of the program was then modified to report LCP values, allowing quantitative analysis of the effect of varying patterning parameters (L, Smax, cL and cR) on LCP values following CO designation. As an example, the effect of CO designation patterning on LCP values when L=0.1, Smax=2 and the bivalent has fully clamped ends (cL and cR both equal 1) is shown in Fig. 6.

**Fig. 6.**
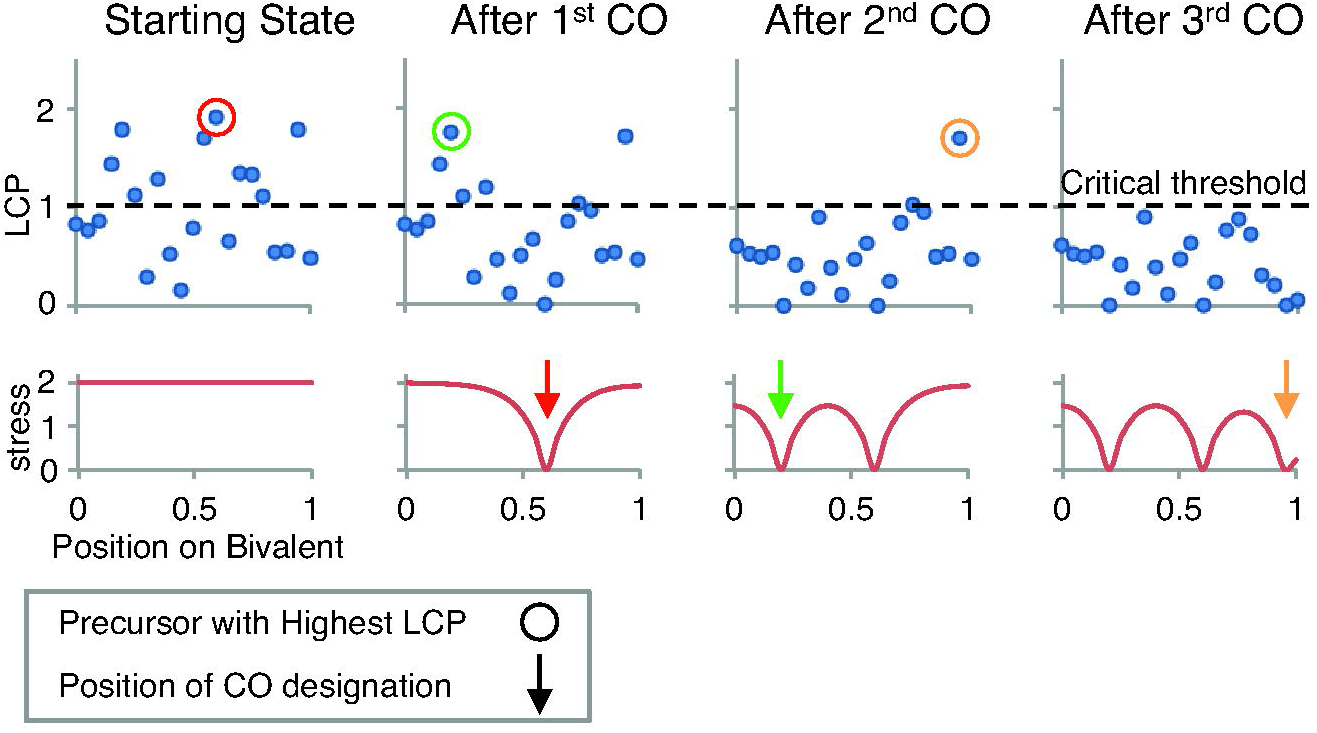
A quantitative example of crossover designation patterning along a single bivalent by the Beam-Film model. 21 precursors (N=21) were evenly positioned (E=1) along a single (n=1) bivalent (at 0, 0.05, 0.1…0.95, 1) and assigned intrinsic sensitivities in accord with parameter A=1. The maximum stress level (Smax) was set to 2, the stress relief distance (L) was set to 0.1 (fraction of bivalent length) and the ends were fully clamped (cL=1, cR=1). The local crossover potential (LCP) of each precursor (top row, blue dots) and the level of stress along the bivalent (bottom row, red line) were then calculated, first prior to any CO designations (“Starting State”) and then as the CO designation process progresses to completion. The LCP is the product of its intrinsic sensitivity and the local level of stress at the corresponding position (text). The “critical threshold” is the minimum value of the LCP at which a CO-designation can occur. This analysis was achieved by modifying the pattern_event_designations_according_to_beam_film_model function of the program to report LCP values for each precursor at each stage. The stress at each precursor position was calculated by dividing the LCP values for each precursor by its intrinsic sensitivity. When CO designation initiates, the precursor with the highest LCP (red circle) gives rise to the first CO designation (red arrow). Resulting local stress relief and redistribution produces a change in the level of stress along the bivalent (red line). This, in turn, changes the LCPs for all remaining precursors (compare positions of blue dots before and after the first CO designation). The same sequence of events occurs two more times, giving rise to a second and third CO designation (green and orange circles/arrows). At this point, none of the remaining precursors has an LCP above the critical threshold and the process stops, resulting in three crossover designations at positions 0.2, 0.6 and 0.95.

### 3.5 How to Make Files Accessible to MATLAB

1. Open MATLAB.
2. Add the files contained in the folder Crossover_Simulation_and_Analysis to the MATLAB path. The simplest way to do this is to use the navigation bar at the top of the screen to open the unzipped folder Crossover_Simulation_and_Analysis (*see* **Note 3**).

### 3.6 How to Run a Simulation

1. Create a table (*e.g.* using the program Excel) listing the parameters to be run in the format indicated in Table 1. The table should have headers (*see* **Note 4**). Each parameter should be in a separate column, and separate combinations of parameters should be entered on separate rows. An example input table (simulation_parameters.csv) is included alongside the program in the Crossover_Simulation_and_Analysis folder.
2. Save the table as a comma separated values (.csv) file (*see* **Note 5**) into the Crossover_Simulation_and_Analysis folder (*see* **Note 6**).
3. Type the following into the Command Window: ≫crossover_simulation(‘file_name.csv’) where file_name.csv is the name of the file containing the input table (see **Notes 7** and **8**).
4. Press return/enter
5. The program will simulate crossover patterning for each combination of parameters in the input table (*see* **Note 9**)
6. The results of the simulation will be saved in the current folder (Crossover_Simulation_and_Analysis) and take the form of .csv files named file_name.csv_lineX.csv, where file_name.csv is the name of the file containing the input table and X is the row number containing the parameters set used for the simulation (*see* **Note 10**). A separate file is saved for each parameter set.

### 3.7 How to Analyze a Set of Real or Simulated Crossover Positions

1. Save a copy of the crossover patterns (real or simulated) to be analyzed, into the Crossover_Simulation_and_Analysis folder, *see* **Note 6**. The data should be in the format of a table (without headers). The bivalent length should be entered in column 1, with crossover positions entered in subsequent columns. Values for different bivalents should be entered on different rows. The simulation function of the program automatically produces a table in the correct format (and saved to the correct folder). It therefore does not need to be modified for analysis.
2. Type the following into the command window: ≫analyze_events_on_linear_objects(‘file_name.csv’,[]) where file_name.csv is the name of the file containing the crossover positions for analysis (*see* **Note 7** and **Note 11**).
3. Press return/enter.
4. The program will analyze the crossover positions and calculate the coefficient of coincidences using the default number of intervals (*see* **Note 11**).
5. The program will save the results in the form of a .csv file named ‘file_name.csv_Xintervals.csv’, where file_name.csv is the name of the parent file that was analyzed and X is the number of intervals used for CoC analysis. The format of the output table is shown in Table 2 (*see* **Note 12**).

**Table 2.** Output of the function for analysis of experimental or simulated data sets. The analysis function of the program produces results in the format of a table with 15 columns (complete with headers), saved as a comma separated values (.csv) file. The number of rows will vary depending on the data being analyzed and with large datasets may exceed the maximum number of rows that some software (such as Microsoft Excel) can open. If this occurs, in order for users to view the complete list of distances between adjacent crossovers (the column that may exceed the maximum number of rows), users can use software such as R or MATLAB. The values in the table can be used to generate graphs using a number of different available software. Note that MATLAB will fill empty cells of the table with a value of ‘NaN’. These values can be removed, for example, by using the find and replace function of Excel. ^a^For simulations, this will always equal 1. ^b^Units are equal to the units of measurement in the input table (*e.g.* μm or Mbp).

| Column |  | Header | Description/Contents |
| --- | --- | --- | --- |
| 1 | A | Number of analyzed objects | Total number of measured/simulated bivalents (= “n” for a simulation output) |
| 2 | B | Mean object length | The mean length of all measured<sup>a</sup> bivalents<sup>b</sup> |
| 3 | C | Mean number of events per object | The average number of total crossovers per bivalent |
| *Distribution of [total crossovers per bivalent] (e.g. see Fig. 5A)* |  |  |  |
| 4 | D | Total number of events | The range of total crossovers per bivalent observed in the measured population |
| 5 | E | Normalized frequency | The proportion of bivalents with the corresponding total number of crossovers defined by column 4/D |
| *Bivalent interval analysis (e.g. see Fig. 5B and 5C)* |  |  |  |
| 6 | F | Interval | The center position of all intervals into which the bivalent has been divided<sup>b</sup> |
| 7 | G | Frequency of events | The frequency with which a crossover occurred within each of the intervals defined by column 6/F |
| 8 | H | Inter-interval distance | The center-to-center distances between all possible pairs of intervals used for Coefficient of Coincidence (CoC) analysis<sup>b</sup> |
| 9 | I | Mean CoC | The mean CoC corresponding to the inter-interval distance defined by column 8/H |
| *Analyses of distances between adjacent crossovers* |  |  |  |
| 10 | J | Mean distance between adjacent events | The average distance between adjacent crossovers<sup>b</sup> |
| **11** | **K** | Distance between adjacent events | A list of all the measured distances between adjacent crossovers<sup>b</sup> |
| **12** | **L** | Gamma shape MLE | The maximum likelihood estimate of the shape value of the best fit gamma distribution for the measured distances between adjacent crossovers (listed in 11/K) |
| **13** | **M** | Gamma shape CIs | 95% confidence intervals of the value in 12/L |
| **14** | **N** | Gamma scale MLE | The MLE of the scale value of the best fit gamma distribution for the measured distances between adjacent crossovers (listed in 11/K) |
| **15** | **O** | Gamma scale CIs | 95% confidence intervals of the value in 14/N |

## 4. Notes

1. The program ‘Crossover Patterning Simulation and Analysis 1.0’ is composed of 16 MATLAB files (for names *see* **Note 3**) and is available for download *via* a link on the authors’ website http://projects.iq.harvard.edu/kleckner_lab. Please contact the authors if the weblink is not available. The files are contained in the folder ‘Crossover_Simulation_and_Analysis’, which must be unzipped before use. Users can rename this folder as they see fit. We recommend that users save a copy of the ‘Crossover_Simulation_and_Analysis’ folder to the ‘MATLAB’ folder on their computer to simplify adding the files to the file path (see Subheading 3.5, **step 2**).
2. The program has been tested on MATLAB versions R2014b and R2015a on both Windows and MAC OS environments.
3. The following MATLAB files should be visible in the ‘Current Folder’ panel (typically found on the left-hand side of the screen); add_typeII_events.m, analyze_events_on_linear_objects.m, conc.m, crossover_simulation.m, distribute_precursors_evenly_along_objects.m, event_per_object_from_population_mean.m, event_spacing.m, extract_information.m, generate_precursor_array.m, generate_precursor_sensitivities.m, generate_report_table.m, interval_analysis.m, mature_designated_precursors.m, padcat.m, pattern_event_designations_according_to_beam_film_model.m, and summarystatistics.m.
4. The program ignores the first line of the table as being headers. The user can therefore label the parameters as they see fit. However, it is necessary to keep the order of parameters as described in Table 1.
5. The program is capable of correctly recognizing other file types (*e.g.* .xlsx). For more information, search MATLAB help for information on its readtable function.
6. It is not strictly necessary to save either the file containing the input table for simulation or the list of CO positions for analysis to the folder containing the program functions. In general, users that are familiar with MATLAB will be able to depart from the described method at numerous stages.
7. The symbol ‘≫’ indicates the MATLAB command line. The user should not enter this part of the command (it should be present automatically). The user should be careful if entering commands by copying and pasting. MATLAB tends to have a problem recognizing the ‘’ symbols from other programs/file types. In MATLAB R2015a, the user will know if the quotation marks have been recognized as they will turn purple (along with the file name). MATLAB commands are case sensitive. The user must enter the command and corresponding file name accurately (*e.g.* do not add spaces).
8. The length of time required to run a simulation depends on a number of factors, but scales approximately with the number of precursors in the simulation (no matter how they are divided between bivalents and separate simulations). For reference, the author finds that it takes approximately 2s for every 10,000 precursors total (average number of precursors per bivalent*number of simulated bivalents*number of parameter sets), on a standard MacBook Pro laptop.
9. The program runs a separate simulation for each parameter set *(i.e.* row of the input table) sequentially. Therefore, if a parameter set were to cause the program to crash, it will not move on to the next parameter set.
10. Output file size will depend on the simulation, but is unlikely to be larger than 1 Mb for a simulation of 10,000 bivalents.
11. As a default the program divides each chromosome into a number of intervals that is equal to 1/(the mean inter-crossover distance)*5. The user can input a defined number of intervals. For example, the command ≫analyze_events_on_linear_objects(‘file_name.csv’,10) would analyze the crossover positions using 10 intervals for CoC analysis. If users do not want to use intervals of equal length (*e.g.* for analysis of genetic crossovers), they can manually define the boundary of each interval. For example, the command: ≫analyze_events_on_linear_objects(‘file_name.csv’, [0.1,0.2,0.6,0.9]) will analyze crossovers in the following three intervals: 0.1-0.2, 0.2-0.6 and 0.6-0.9. In this case, it is important that the user encloses the list of boundary positions with square brackets and separates each interval boundary with a comma (no space).
12. The output table can be viewed and manipulated using a number of different software including Microsoft Excel. It is important to note that some software (such as Microsoft Excel) have a maximum number of rows that can be imported. It is possible to exceed this limit as large datasets (real or simulated) can produce many calculated distances between adjacent COs. If this occurs, and the user would like to access the full list of distances between adjacent COs, they should open the .csv file using a program that can handle many rows, such as R or MATLAB.

## Acknowledgements

The above methods were developed by the authors in collaboration with Prof. John Hutchinson (SEAS; Harvard University). This work and S.W. were supported by a grant to N.K. from the N.I.H. (2RO1 GM 044794). M. A. W. is supported by HFSP long-term fellowship LT000927/2013. L. Z. is funded by the 1000-talents Plan for young researchers (41200095551503) and by Shandong University (11200085963001). We would like to thank Guillaume Witz and Frederick Chang for help with updating the MATLAB code; and Terry Hassold and Pat Hunt for communication of unpublished Mlh1 focus data for human male chromosomes. This manuscript is currently in press for the forthcoming book Methods in Molecular Biology (Meiosis) http://www.springer.com/us/book/9781493963386.

## References

1. Hunter, N. (2015). Meiotic Recombination: The Essence of Heredity. Cold Spring Harb Perspect Biol 7: a016618

2. Zickler, D., Kleckner, N. (2016). Recombination, pairing and synapsis of homologs during meiosis. Cold Spring Harb Perspect Biol 7: a016626

3. Wang, S., Zickler, D., Kleckner, N. et al. (2015). Meiotic crossover patterns: obligatory crossover, interference and homeostasis in a single process. Cell Cycle 14, 305–314

4. Kleckner, N., Zickler, D., Jones, G. H. et al. (2004). A mechanical basis for chromosome function. Proc Natl Acad Sci USA 101, 12592–12597

5. Zhang, L., Liang, Z., Hutchinson, J. et al. (2014). Crossover patterning by the Beam-Film Model: Analysis and implications. PloS Genetics 10: e1004042

6. Zhang, L., Wang, S., Yin, S. et al. (2014). Topoisomerase II mediates meiotic crossover interference. Nature 511, 551–556

7. Zhang, L., Espagne, E., de Muyt, A. et al. (2014). Interference-mediated synaptonemal complex with embedded crossover designation. Proc Natl Acad Sci USA 111, E5059–E5068

8. Anderson L.K., Lohmiller L.D., Tang X. et al. (2014). Combined fluorescent and electron microscopic imaging unveils the specific properties of two classes of meiotic crossovers. Proc Natl Acad Sci U S A 111, 13415–13420

9. Wu T.C., Lichten M. (1995). Factors that affect the location and frequency of meiosis-induced double-strand breaks in Saccharomyces cerevisiae. Genetics 140, 55–66

10. Garcia, V., Gray, S., Allison, R.M. et al. (2015). Tel1^ATM^-mediated interference suppresses clustered meiotic double-strand-break formation. Nature 520, 114–118

11. Gruhn J.R., Rubio C., Broman K.W. et al. (2013). Cytological studies of human meiosis: sex-specific differences in recombination originate at, or prior to, establishment of double-strand breaks. PloS One 8: e85075

